# Exploring the role of cellular homologous of the 30K-superfamily of plant virus movement proteins

**DOI:** 10.1101/245472

**Authors:** José L. Carrasco, Jesús A. Sánchez-Navarro, Santiago F. Elena

**Author notes:** Equal contribution (José L. Carrasco, Jesús A. Sánchez-Navarro). Corresponding author: *E-mail address* (S.F. Elena).

## Abstract

Genes orthologous to the 30K-superfamily of movement proteins (MP) from plant viruses have been recently discovered by bioinformatics analyses as integrated elements in the genome of most vascular plants. However, their functional relevance for plants is still unclear. Here, we undertake some preliminary steps into the functional characterization of one of these putative *MP* genes found in *Arabidopsis thaliana*. We found that the *AtMP* gene is expressed at different stages of the plant development, with accumulation being highest in flowers but lowest in mature siliques. We also found down-regulation of the gene may result in a small delay in plant development and in an exacerbation of the negative effect of salinity in germination efficiency. We have also explored whether changes in expression of the endogenous *AtMP* have any effect on susceptibility to infection with several viruses, and found that the infectivity of tobacco rattle tobravirus was strongly dependent on the expression of the endogenous *AtMP*. Finally, we have cloned the endogenous *MP* from four different plant species into an expression vector that allows for specifically assessing their activity as cell-to-cell movement proteins and have shown that though some may still retain the ancestral activity, they do so in a quite inefficient manner, thus suggesting they have acquired a novel function during adaptation to the host genome.

## 1. Introduction

Endogenous viral elements (EVE) are widespread in eukaryotic genomes. EVEs result from the integration of a dsDNA copy of a portion of viral genomes into the chromosomes of the host germinal cell line, thus allowing for its vertical transmission. When first discovered, EVEs were thought to be limited to retroviruses, as this class of viruses are the only known eukaryotic viruses that require integration in the host cell genome to complete their lifecycle and encode the necessary molecular machinery, the reverse transcriptase (*e.g*., Harper et al., 2002). In the last decade or so, however, plenty of examples have been published of EVEs from all major classes of eukaryotic viruses (Holmes, 2011; Feschotte and Gilbert, 2012; Aiewsakun and Katzourakis, 2015). Indeed, non-retroviral integrated RNA virus sequences (NIRV; Koonin, 2010) are a particular class of EVEs whose closest free-living counterparts are *bona fide* RNA viruses of both positive and negative senses. The mechanism by which exogenous RNA viruses can be endogenized is not fully understood yet, but some evidences point towards the involvement of the enzymatic machinery encoded by the retrotransposons residing in the host genome (Geuking et al., 2009; Taylor and Bruenn, 2009; Horie et al., 2010).

EVEs provide the raw material for evolutionary exaptations, that is a shift in the function of a trait. Well-known examples of such exaptations are the mammalian syncytin-A endogenized from a retrovirus *env* gene necessary for the morphogenesis of placenta (Mi et al., 2000), or the polydnaviruses endogenized by braconid and ichneumoid wasps that act as virulence factors that allow them to parasitize the larvae of their insect hosts (Drezen et al., 2017). Some less known but highly interesting examples of exaptations have also been described for plant NIRVs. For example, Tanne and Sela (2005) described the integration of coat protein (CP) from potato potyvirus Y (PVY) into the genome of grapevine via a process of nonhomologous recombination between the viral genomic RNA and a host class II retrotransposon. The *CP* gene expression was confirmed by several techniques. It was later hypothesized that such gene expression may provide resistance to infection by other related viruses with sequence homology to the endogenous copy (Bertsch et al., 2009). In a second remarkable example, Chiba et al. (2011) found high sequence identity between the product from gene *ILR2* of *Arabidopsis thaliana* and the *CP* gene of *Rosellinia necatrix* partitivirus 2. *IRL2* regulates the expression of auxin and thus is essential for the coordinated growth of plants in response to environmental stresses, suggesting that the viral CP acquired a completely novel function here. In a final example, the protein encoded by the *gem* gene of the meadow fescue *Festuca pratensis*, involved in leaf senescence, shares sequence identity to the CP of *Sclerotinia sclerotionum* partitivirus S (Liu et al., 2010).

The most common approach to discover EVEs is by bioinformatics analyses of whole cellular genomes and transcriptomes that are publicly available (Bertsch et al., 2009; Liu et al., 2010; Chiba et al., 2011; Cui and Holmes, 2012; Geering et al., 2014; Mushegian and Elena, 2015). Unfortunately, the cellular functions of these NIRVs remain unclear, owed to a lack of experimental work to confirm the *in silico* findings. Recently, Mushegian and Elena (2015) performed such an *in silico* study focusing on a particularly widespread gene among different families of plant RNA viruses and pararetroviruses, the 30K-superfamily of movement proteins (MP). Comparative analyses of different 30K MPs have revealed many properties of relevance to the mechanism of cell-to-cell spread of plant viruses (Mushegian and Elena, 2015; and references therein): (*i*) the ability to bind ssRNA and DNA cooperatively but non-specifically, (*ii*) their localization in the plasmodesmata and with the endoplasmic reticulum or with actin cytoskeleton that scaffolds the reticulum and links it to the plasmodesmata, (*iii*) the capacity to increase the plasmodesmatal size exclusion limit, (*iv*) self-association *in vivo* to form hollow tubular structures, and (*v*) MPs have been shown to interact with diverse host plant proteins, some of which are of functional significance. Mushegian and Elena (2015) found that 30K MPs from the *Caulimoviridae* were present in all euphyllophyte plants (all vascular plants except mosses), with many instances retaining uninterrupted ORFs that appear to be under positive selection at the codon level, suggesting that they must be playing a molecular function in the cell. Interestingly, a putative *MP* sequence was found next to predicted gene *At1g37113* in *A. thaliana*. These observations prompted the authors to suggest that the expression of these endogenous *MP*s may have impacted the interaction between plants and viruses.

Here, we sought to explore the above hypothesis. To do so, we have performed two classes of exploratory experiments. Firstly, for practical reasons, we have focused on the analysis of the *A. thaliana MP*-*At1g37113* gene fusion, hereafter referred as *AtMP*. We have first confirmed the expression of *AtMP* by RT-qPCR in different tissues and developmental stages. Next, we generated transgenic plant lines expressing artificial microRNAs (amiRs) against the putative *AtMP* that knock-down its expression. We sought for phenotypes associated to these knock-down mutations as well as to their effect on the infectivity of several viruses. Secondly, we have cloned cDNAs for a number of putative *MP*s from different plant species detected by Mushegian and Elena (2015) into an alfalfa mosaic alfamovirus (AMV)-based expression vector that allows to specifically test the activity of any given protein of the 30K-superfamily as a movement one.

## 2. Materials and methods

### 2.1. Plant material and growth conditions

*A. thaliana* ecotype Col-0 was used in the study of *MP*-*At1g37113* expression. Plants were grown under different photoperiod conditions depending on the experiment. Long-day photoperiod (16 h light, 25 °C/8 h dark, 20°C) was used for plant transformation and tissue expression analysis. For virus inoculation, plants were grown under short-day photoperiod (8 h light, 25 °C/16 h dark, 20 °C).

To analyze the germination efficiency of the different *A. thaliana* genotypes used in this study, seeds were sterilized by washing them with 70% ethanol supplemented with 0.05% SDS for 3 min, followed by 96% ethanol for 1 min. Then they were allowed to dry on a piece of filter paper and transferred to MS medium plates with the help of a sterile toothpick. To explore the effect of salinity on germination efficiency, MS plates where supplemented with 0, 100 or 150 mM NaCl. Germination rates were calculated using the Maguire’s index (McNair et al., 2012).

### 2.2. Design and construction of amiR-expressing transgenes

For the purpose of down-regulating the expression of *AtMP* gene in Arabidopsis, an amiR-based strategy was followed (Schwab et al., 2006). Overexpression in transgenic plants of constructs generating amiRs specifically targeted against the *AtMP* might eventually lead to a significant reduction in the expression of that gene. For the design and cloning of amiRs, tools and protocols available in the website MicroRNA Designer application (wmd3.weigelworld.org; Ossowski et al., 2008) were used. The search for specific amiRs was performed selecting default settings and the TAIR10 transcript library (www.arabidopsis.org). Among the possible amiRs provided by the application, three were finally selected on the basis of the criteria previously established by Schwab et al. (2005, 2006) (Table 1).

**Table 1.** AmiRs selected for generation of transgenic lines.

| **amiR** | **Sequence<sup>a</sup>** |
| --- | --- |
| amiR1 | TTAGGACCGAAT**CtAc**GCCTT |
| amiR2 | AATAGGGACTTTG**tTCTCgC** |
| amiR3 | **t**TTGTTCTAGTTG**Ac**AGGCTA |

<sup>a</sup>Bold lowercase letters correspond to mismatches with respect to the target sequence.

The selected sequences were introduced in the context of the miR319a precursor for subsequent expression in planta. The miR319a backbone is present in the pRS300 plasmid (Schwab et al., 2006). Three rounds of PCR amplification were performed for each amiR using pRS300 as template and six oligonucleotide primers listed in Supplementary Table S1. Four of these primers were specific for each selected amiR (designated I-IV) and two annealed to the pRS300 plasmid (A and B). As a result, for each construct, three partially overlapping fragments were generated, which were then used as template in a final PCR amplification with flanking A and B primers to generate the miR319a-based amiR construct.

With the aim of cloning the amiR constructs, an additional PCR amplification was performed, using primers containing attB sites suitable for Gateway cloning technology (amiRNA-attB1 and amiRNA-attB2; Supplementary Table S1), which anneal to positions internal to those of the A and B primers (Jover-Gil et al., 2004). Each amplified product was cloned into pDONR207 and transferred into the pMDC32 destination vector for overexpression *in planta* (Curtis and Grossniklaus, 2003).

### 2.3. Generation and selection of transgenic plants

Arabidopsis plants were transformed by the *Agrobacterium tumefaciens*-mediated floral dip method (Clough and Bent, 1998), following modifications described in Logemann et al. (2006). Selection of transformants was carried out in MS medium supplemented with hygromycin B according to Harrison et al. (2006). Single-insertion lines were selected for each amiR construct by marker segregation analysis.

### 2.4. Quantification of the endogenous AtMP expression in A. thaliana

RNA was purified from plant tissue with the GeneJet Plant RNA Purification Mini kit (Thermo Scientific), according to instructions provided by the manufacturer. For tissue-specific expression, different sets of plants were grown simultaneously for each tissue sample. For expression in leaves, plants were grown for about 4 weeks (growth stage about 3.70 in the Boyes’ scale; Boyes et al., 2001). Stems, flowers, leaves, and siliques were collected from five plants that were 6 weeks old (growth stage about 6.50) and pooled according to the tissue. Gene expression was analyzed by RT-qPCR in a StepOnePlus Real-Time PCR System (Applied Biosystems) using the GoTaq 1-Step RT-qPCR kit (Promega) following manufacturer’s recommendations (see Supplementary Table S1 for details on the oligonucleotide primers used for amplification). The StepOne Software v.2.2.2 (Applied Biosystems) was used for data analysis according to the comparative *C_T_* method (Schmittgen and Livak, 2008), with *At1g13320* (*PP2A-A3*, subunit A3 of the serine/threonine phosphatase type 2A) and *At2g28390* (*MON1*, a SAND family protein) used as reference genes (Czechowski et al., 2005).

### 2.5. Virus, plant inoculations and verification of infection

The viruses selected to examine the susceptibility to virus infection of amiR transgenic lines as compared to wildtype *A. thaliana* Col-0 plants are listed in Table 2. Infectious crude extracts were prepared by homogenizing material from infected *Nicotiana benthamiana* plants in 10 volumes of inoculation buffer (50 mM potassium phosphate pH 7.0, 3% PEG). Ten five-week-old plants were mechanically inoculated by gently rubbing 5 µl aliquots of infectious crude extract supplemented with 10% Carborundum onto the surface of three different fully expanded rosette leaves. After inoculation, plants were maintained under the same growth conditions (short-day photoperiod).

**Table 2.** Viruses used to analyze the susceptibility of the amiR-expressing lines to infection.

| **Virus** | **Genus (Family)** | **Genome** |
| --- | --- | --- |
| *Alfalfa mosaic virus* (AMV) | *Alfavirus* (*Bromoviridae*) | Tripartite linear ssRNA(+) |
| *Plum pox virus* (PPV) | *Potyvirus* (*Potyviridae*) | Monopartite linear ssRNA(+) |
| *Tobacco mosaic virus* (TMV) | *Tobamovirus* (*Virgaviridae*) | Monopartite linear ssRNA(+) |
| *Tobacco rattle virus* (TRV) | *Tobravirus* (*Virgaviridae*) | Bipartite linear ssRNA(+) |
| *Turnip mosaic virus* (TuMV) | *Potyvirus* (*Potyviridae*) | Monopartite linear ssRNA(+) |

Fourteen days post-inoculation (dpi), the complete rosette of inoculated plants was harvested, frozen and stored at –80°C. Infection with RNA viruses was determined by RT-PCR, with RNA being prepared as already mentioned. Oligonucleotides used as primers in all these reactions are listed in Supplementary Table S1.

### 2.6. Amplification and cloning of 30K-like cellular orthologues in AMV-based expression vector

A modified infectious AMV cDNA 3 clone (Fig. 4), which expresses the green fluorescent protein (GFP) (pGFP/A255/CP) (Sánchez-Navarro and Bol, 2001), was used to exchange the N-terminal 255 amino acids of the AMV *MP* gene with the putative MP gene of *Arabidopsis lyrata* (291 amino acids long; GenBank XM_002893778; GenBank XP_002893824), *A. thaliana* (414 amino acids long; GenBank NM_001123950; GenBank NP_001117422), *Lactuca sativa* (326 amino acids long; GenBank AFSA01069977), or *Nicotiana tabacum* (317 amino acids long; GenBank AJ414165; GenBank CAC88784). The putative endogenous *MP* genes were amplified by using specific primers (Supplementary Table S1). The digested fragments were used to replace the *Nco*I–*Nhe*I fragment of pGFP/A255/CP, corresponding with the N-terminal 255 amino acids of the AMV MP, to generate the constructs pGFP/*A.lyrata*:A44/CP, pGFP/*A.thaliana*:A44/CP, pGFP/*N.tabacum*:A44/CP, and pGFP/*L.sativa*:A44/CP. A negative control corresponding to the *A. thaliana* gene lacking the 5’ terminal 38 nucleotides, which allow to use a start ATG codon located in a different frame (fs), was generated.

### 2.7. Inoculation of P12 plants

pGFP/A255/CP and the corresponding *MP* derivatives, were linearized with *Pst*I and transcribed with T7 RNA polymerase. The transcripts were inoculated onto transgenic *N. tabacum* plants that express constitutively the P1 and P2 polymerase proteins of AMV (P12), as previously described (Taschner *et al*., 1991). The fluorescence derived from the chimeric AMV RNA 3 carrying the GFP, was monitored using a Zeiss LSM 780 confocal microscope at 3 dpi, with excitation at 488 nm and emission at 510–560 nm. A number of fluorescents objects was counted, and classified as single infected cells or as multi-cellular infected foci. The frequency of foci was then estimated using the LaPlace estimator for the Binomial parameter, which produces more robust estimates than the commonly used maximum-likelihood estimator for small sample sizes (Agresti and Coull, 1998).

## 3. Results

### 3.1. *Expression of AtMP varies among tissues in* A. thaliana

First, we sought to prove whether the predicted *A. thaliana* endogenous *AtMP* gene was expressed in different plant tissues. To do so, we sampled leaves, stems, flowers and siliques from 5 plants and evaluated the expression of the *AtMP* gene by RT-qPCR relative to two reference genes that have been shown to express in a highly reproducible and constant manner across different tissues and under different stresses, *PP2A-A3* and *MON1*. Fig. 1 shows the results of these expression analyses. A first consideration has to be made about the differences in expression depending on which reference gene is used. In general, relative expression values are consistently smaller for *PP2A-A3* than for *MON1*, however, the rank order of tissues is highly consistent among both reference genes: highest expression occurs in flower (∼2–3 times more than in leaves), whereas in siliques expression it is almost null (∼26–74 times less than in leaves). Next, we sought to evaluate whether these apparent differences are actually supported statistically. Oneway ANOVA tests run for both expression datasets found a highly significant difference among tissues (*F*_3,8_ ≥ 452.948, *P* < 0.001 for both reference genes). Notice that data were first log-transformed to ensure homocedasticity of variances among tissues.

**Fig. 1.**
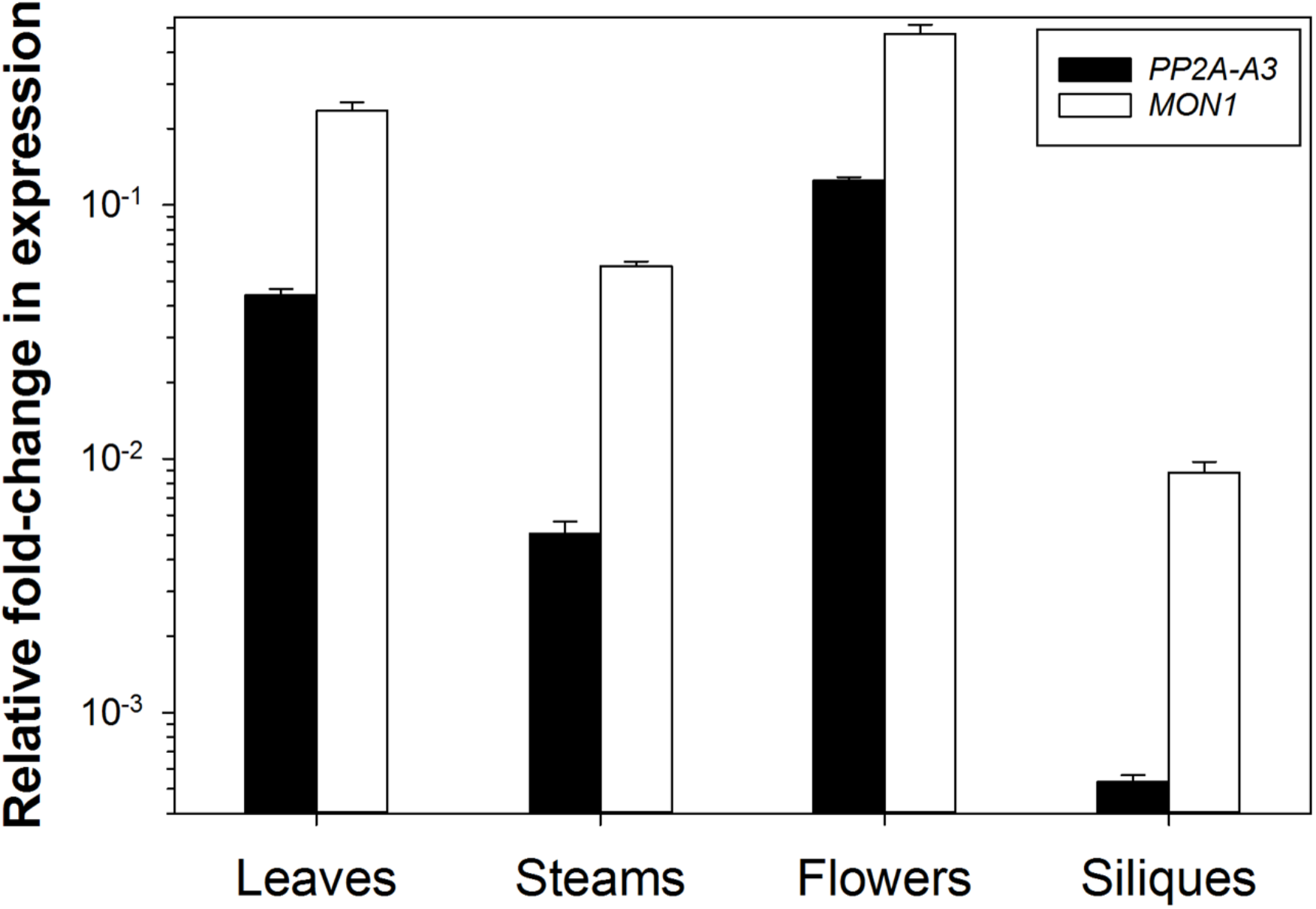
Relative expression of *AtMP* in different tissues of *A. thaliana*. Gene expression was measured by relative RT-qPCR against two reference genes (*PP2A-A3* and *MON1*) as described in section 2.4. Results are consistent for both reference genes (Pearson’s *r* = 0.990, 2 d.f., *P* = 0.010). Notice that the *y*-axis is in logarithmic units. Error bars represent ±1 SD and are based in three biological replicates.

Therefore, we conclude that the Arabidopsis endogenous 30K-like *AtMP* gene is expressed in a tissue-dependent manner.

### 3.2. Knock-down of AtMP results in a small developmental delay

Six Arabidopsis transgenic lines, two for each of the amiRs designed were generated. The three lines showing the largest reduction in expression of the *AtMP* gene were selected for the experiments described in this and the in following sections. Using as reference for the RT-qPCR quantifications the *PP2A-A3* gene, line 1.13.1 expresses the amiR1 (Table 1) and shows 90.63% ±0.81 (±1 SEM) reduction in the expression of the endogenous *AtMP* gene. Line 2.2.2, which expresses the amiR2 (Table 1), shows a 57.74% ±1.56 reduction. Line 3.3.4, expressing the amiR3 (Table 1), shows the smallest, yet significant, 41.89% ±1.45, down effect in expression. The values obtained using the *MON1* gene as reference are highly similar and not reported here. Differences between the three lines are statistically significant (*F*_2,6_ ≥ 452.948, *P* < 0.001; for both reference genes).

To evaluate the possible effect of knock-down *AtMP* expression in plant development, we focused in two specific stages of Boyes’ scale: stage 1 that describes early leaf production and rosette growth (between days 8 – 30 post-germination) and stage 8 that focuses on the timing at which siliques ripen. Fig. 2a shows the average values of Boyes’ scale (here reported as 10× the original scale values) for the wildtype and the three knockdown transgenic lines at three different time points during rosette development. Data were fitted to a repeated-measures ANOVA. A significant effect is associated with the repeated measures of each individual plant at increasing days post-germination, which simply reflects plant growth and development during the time frame of our observations (*F*2,142 = 3027.864, *P* < 0.001). More interestingly, significant differences exist among the four genotypes in their development (*F*_3,71_ = 14.466, *P* < 0.001). A *post hoc* Tukey test shows that wildtype plants (average 10.39) are always bigger and more developed than the knock-down plants, with lines 1.13.1 and 3.3.4 having an intermediate development (average 9.83) and Line 2.2.2 showing the largest retard in development (average 9.25). Interestingly, a significant genotype-by-days interaction has been observed (*F*6,142 = 2.845, *P* = 0.012), suggesting that development is not parallel for all genotypes. Indeed, as it can be appreciated in Fig. 2a, regression lines for Lines 1.13.1 and 3.3.4 cross each other between 21 and 25 days post-germination. Furthermore, 25 days post-germination, 9 out of 17 wildtype plants had a visible floral stem, while among the knock-down plants only 2 out of 19 from Line 3.3.4 had already developed it.

**Fig. 2.**
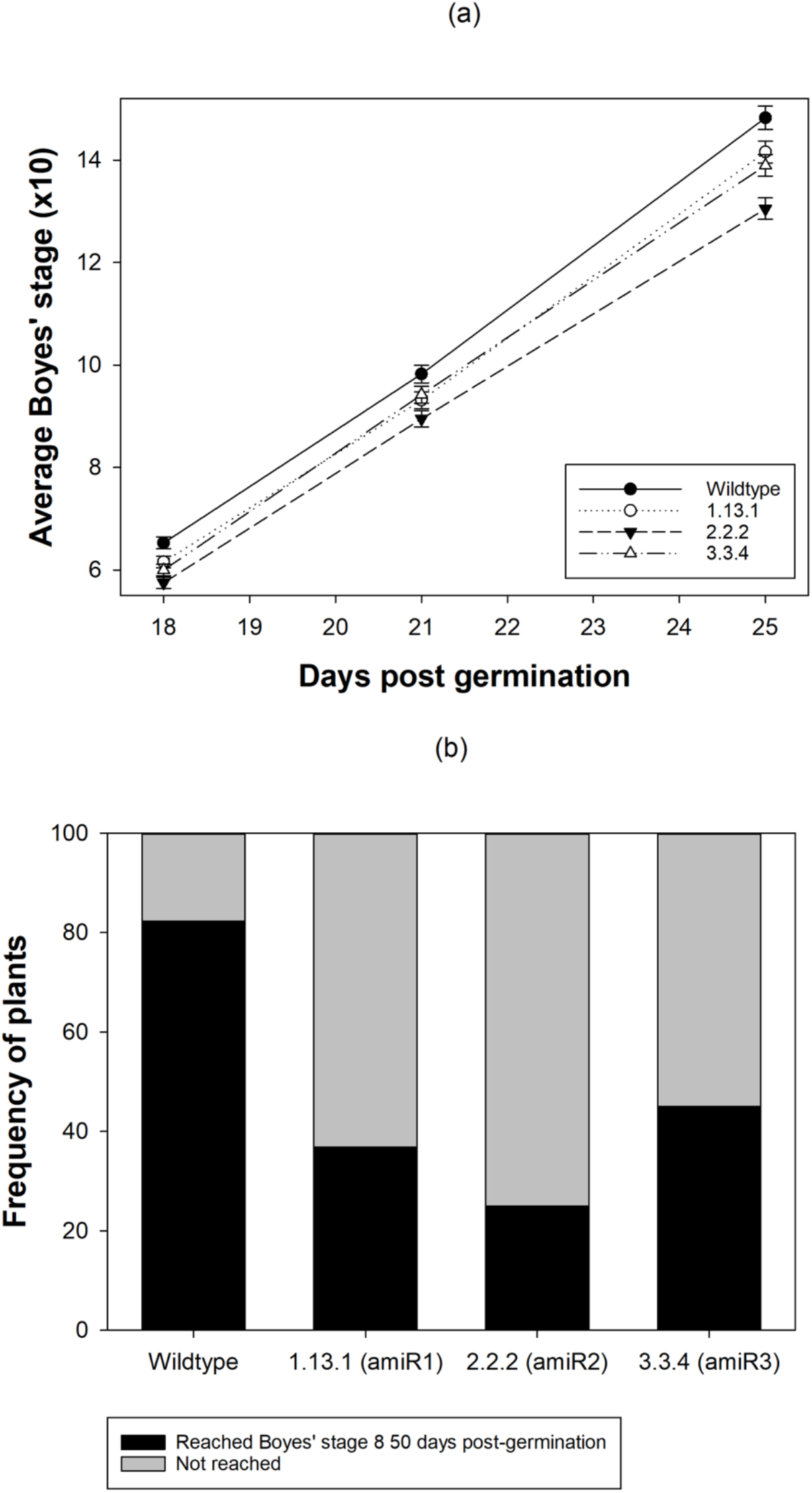
Effect of knocking-down the expression of the *AtMP* gene in plant development. (a) Average Boyes’ scores at 18, 21 and 25 days post-germination for wildtype and the three knock-down amiR lines during stage 1 (leaf production and rosette growth). Error bars represent ±1 SEM and are based in 10 plants per genotype (*n* = 17 for wildtype, 19 for 1.13.1, 20 for 2.2.2, and 19 for 3.3.4). (b) Frequency of plants that reach the silique ripening stage (Boyes’s 8) 50 days post-germination.

Fig. 2b shows the percentage of plants that have reached Boyes’ stage 8 at 50 days post-germination. Overall, significant differences exist among the four genotypes (homogeneity test: χ^2^ = 13.242, 3 d.f., *P* = 0.004), though this difference is entirely driven by the much larger number of wildtype plants that have already ripened at this point in time (82.35%) compared with the three transgenic lines (in the range 25 – 45%).

In conclusion, knocking-down the expression of gene *AtMP* results in a small yet quantitatively significant delay in plant development, which is consistent along the entire life cycle of the plants. However, the magnitude of delay is not obviously correlated to the amount of reduction in *AtMP* expression.

### 3.3. Knock down of AtMP results in reduced germination rate and increased sensitivity to salinity

Seeds from wildtype and the three amiR-expressing lines were placed in MS medium plates supplemented with increasing concentrations of NaCl (ca. 50 seeds per plate, 3 plates per genotype and treatment). Fig. 3a shows the evolution of germination frequencies. Data were fitted to a binary logistic regression equation by means of a generalized linear mixed model (GLMM) approach with a Binomial distribution and a probit link function. The model incorporates days of genotype, days of treatment and NaCl concentration, and all their interactions, as fixed factors and replicate plates as a random factor nested within the interaction between genotype and NaCl concentration. Overall differences exist among the four plant genotypes (χ^2^ = 23.303, 3 d.f., *P* < 0.001), with the largest difference with respect to wildtype being observed for line 1.13.1 (*post hoc* Tukey test *P* < 0.001), and the smallest (yet significant) for line 2.2.2 (*post ho*c Tukey test *P* = 0.032). Increasing NaCl concentration reduces the germination frequency for all genotypes (χ^2^ = 477.870, 2 d.f., *P* < 0.001) but, more interestingly, the effect on germination depends on the plant genotype (χ^2^ = 29.294, 6 d.f., *P* < 0.001). This effect is better illustrated in Fig. 3b, where the rate of germination is shown as a function of salt concentration: compared to the wildtype plants, the rate of germination is strongly affected in lines 1.13.1 and 3.3.4 than in line 2.2.2, with the effect being proportionally more intense as NaCl concentration increased.

**Fig. 3.**
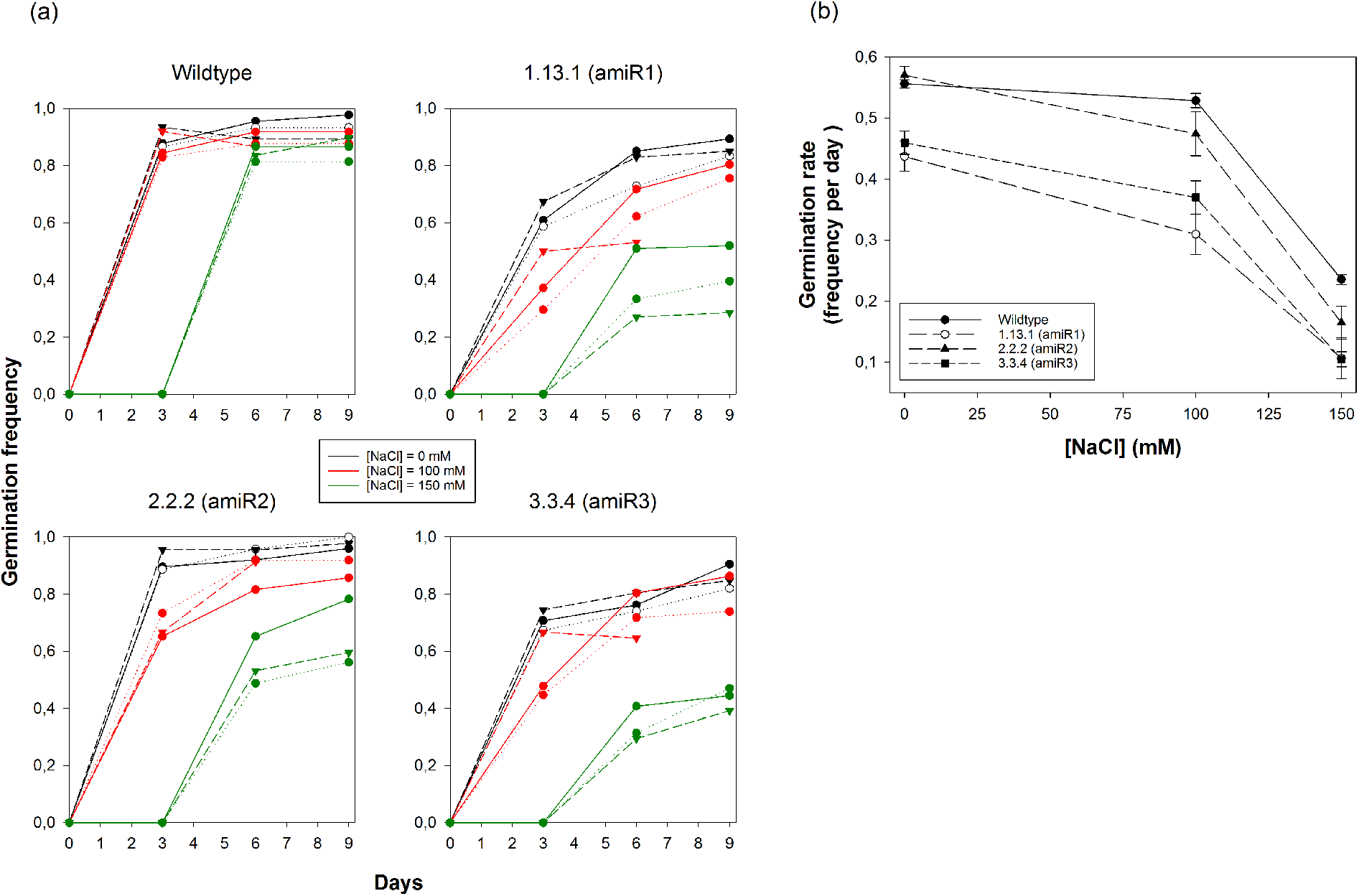
Effect of knocking-down the expression of the *AtMP* gene in germination efficiency at increasing salt concentration. (a) Evolution of germination frequency for the wildtype and the three amiR-expressing lines at the indicated concentrations of NaCl. Different symbols and lines represent the three experimental replicates. (b) Effect of increasing salinity on the germination rate (estimated as the Maguire’s index) for the three amiR-expressing lines and the wildtype.

In conclusion, we show that knocking-down the expression of *AtMP* results in reduced germination efficiency and a stronger effect of salinity in germination rate. Interestingly, the effect is stronger for transgenic line 1.13.1, the one showing the largest reduction in *AtMP* than in transgenic line 2.2.2, that shows a more moderate reduction in the expression of the putative endogenous 30K-like MP.

### 3.4 Knock-down of AtMP affects the infectivity and accumulation of some viruses

To explore whether *A. thaliana* putative endogenous 30K-like MP plays some role during viral infections, we have inoculated wildtype plants and plants expressing amiR1 and amiR2 with a set of RNA viruses varying in their taxonomic degree of relatedness and genome architecture (Table 2). Fig. 4 illustrates the results of these experiments. Infectivity data were fitted to a binary logistic regression equation by means of a GLMM approach with a Binomial distribution and, in this case, a logit link function. Virus species and plant genotypes were treated as orthogonal factors. Overall differences exist among viruses in infectivity (χ^2^ = 54.414, 4 d.f., *P* < 0.001), with the two potyviruses showing the lowest value among all three plant genotypes and AMV and tobacco mosaic tobamovirus (TMV) showing 100% infectivity in all. More interestingly, a significant interaction exists between virus species and plant genotypes, mainly driven by the strong effect that the knock-down in expression of *AtMP* has on the infectivity of tobacco rattle tobravirus (TRV): 33.33% reduction for the amiR1-expresing plants (*post hoc* Tukey test: *P* = 0.049) and 44.44% reduction for the amiR2-expressing plants (*post hoc* Tukey test: *P* = 0.015).

**Fig. 4.**
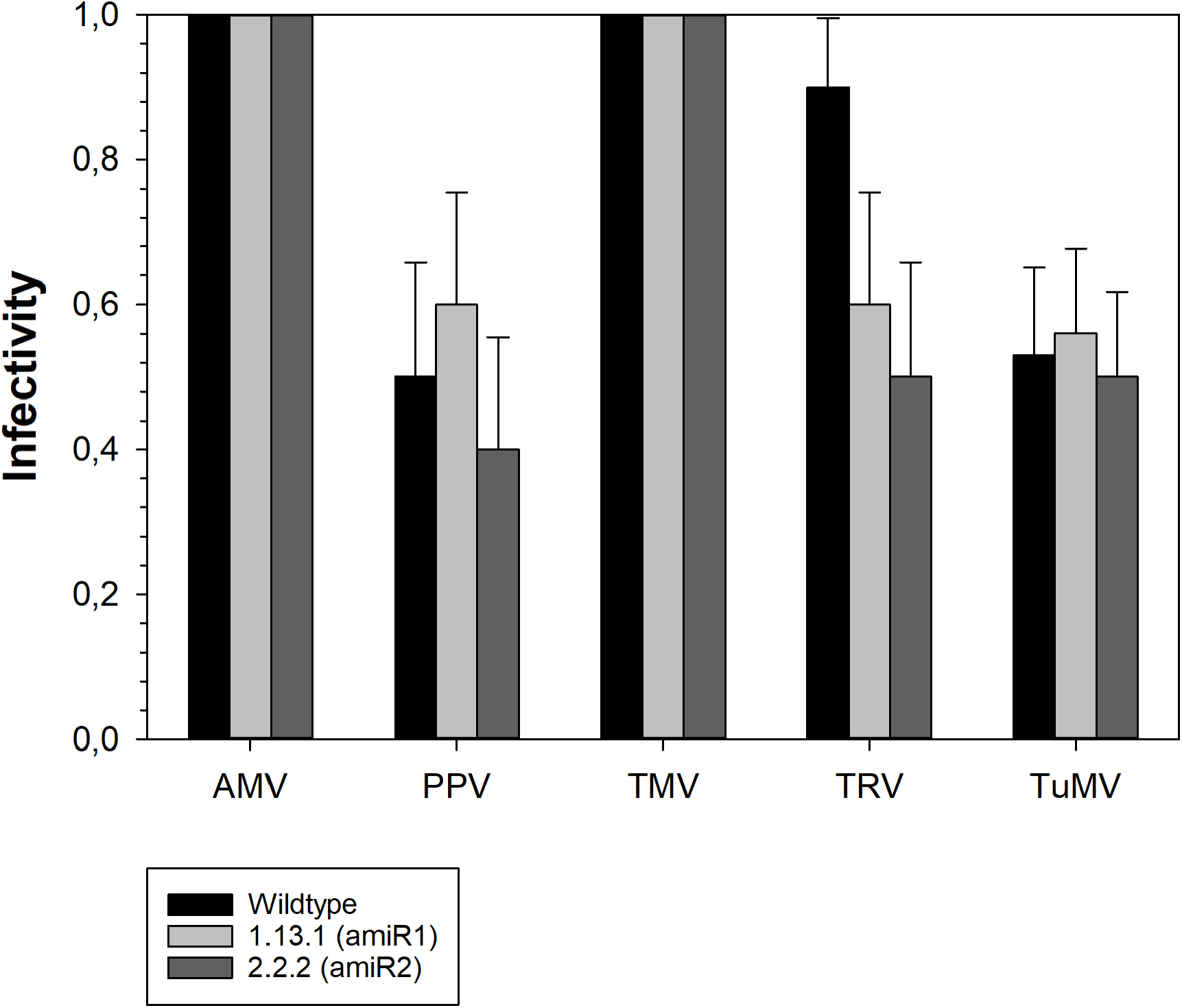
Effect of reducing the expression of the endogenous 30K-like protein in the infectivity of the viruses listed in Table 2. Infectious status was confirmed by RT-PCR using the corresponding pairs of primers listed in Table S1. Infectivity was estimated as the number of plants infected out of 10 plants inoculated. Error bars represent ±1 SE of the estimate of the Binomial parameter.

In conclusion, expression of *AtMP* may have a positive effect in the infectivity of TRV but not in other viruses tested in this exploratory work. By knocking-down the expression of the gene, TRV infectivity is significantly reduced.

### 3.5. Endogenous MPs are still able of moving AMV out of the first infected cell

To extend these results to other putative endogenous *MP* genes found by Mushegian and Elena (2015) in different plant species, we selected those from *A. lyrata, A. thaliana, L. sativa*, and *N. tabacum* based on homology with CaMV sequence, length and availability of material to work with. These genes were cloned in the AMV RNA3-derived expression system for testing their ability to move GFP out of inoculated single cells in *N. tabacum* plants expressing AMV RNAs 1 and 2. Fig. 5 shows a schematic representation of the different clones constructed. For each clone, inoculated leaves were observed 3 dpi with a confocal microscope. The percentages of infectious foci formed by multiple cells are indicated in Fig. 5. The lowest percentage was observed for the *A. thaliana AtMP* gene (14.08%), whereas the larger one was for the *L. sativa* gene (47.92%). In any case, these percentages are significantly smaller than observed for the native AMV *MP* gene (100%). A second clear difference between foci generated by the wildtype AMV and the endogenous *MP*s is the size of the foci. Whereas the average focus generated by wildtype AMV 3 dpi has a mean radius of 10 cells, foci generated by the constructs containing the four endogenous *MP*s were of 3–4 cells radius, on average (Fig. 5). These results, however, have to be taken with some caution, as a few foci (7.69%) of more-than-one cell and of similar morphology were also observed in the control experiment in which a frame shift was introduced into the *A. thaliana AtMP* gene (Fig. 5). Taking this frequency as a proxy to the rate of false positives in this experimental system, the actual values observed for *A. thaliana* and *A. lyrata* cannot be distinguished from experimental noise (Fisher exact tests: *P* ≥ 0.218). However, both the endogenous MPs from *N. tabacum* and *L. sativa* show a significant enrichment in the number of infectious foci formed by many cells (Fisher exact tests: *P* ≤ 0.002 in both cases)

**Fig. 5.**
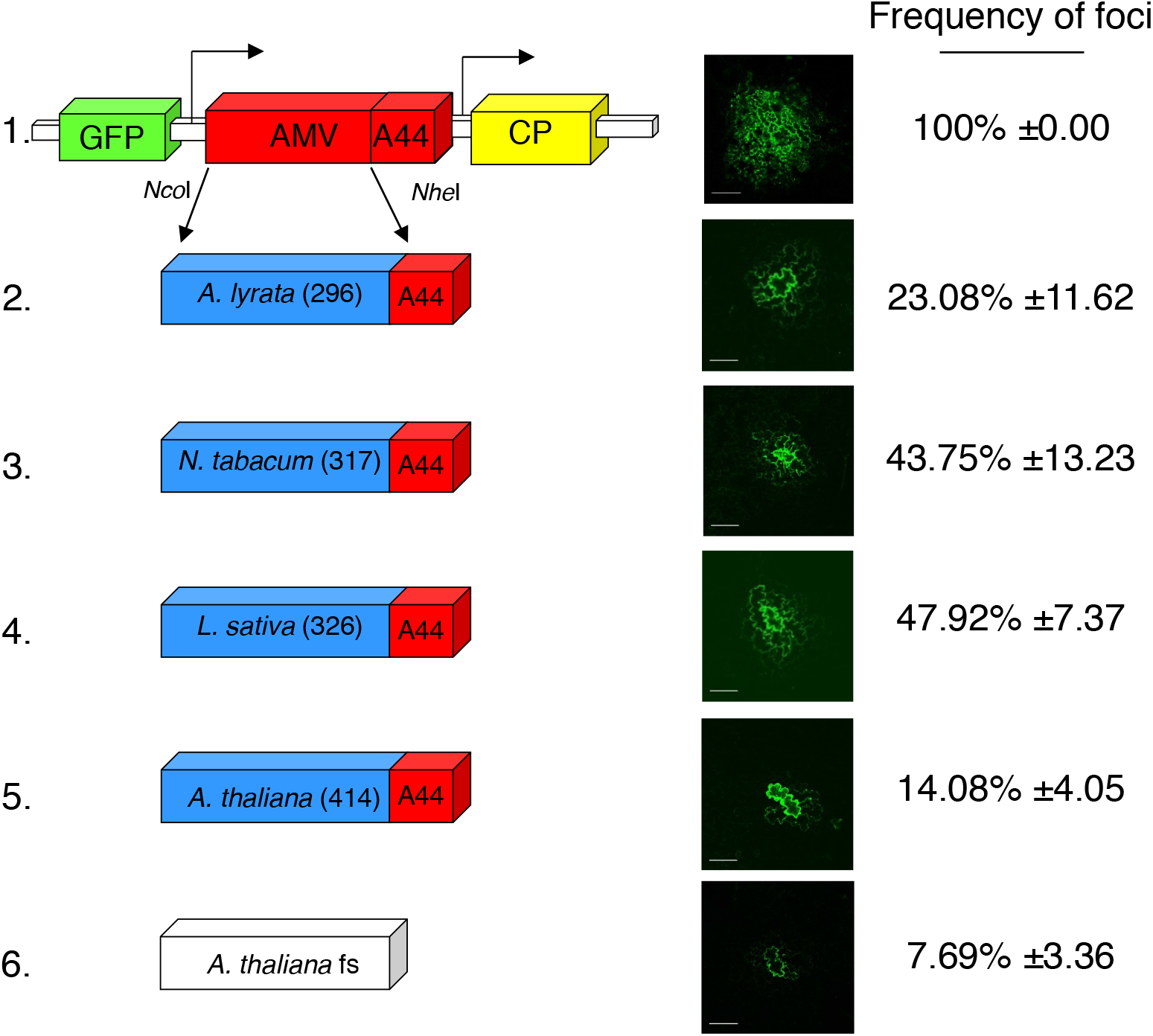
Analysis of the cell-to-cell transport of hybrid AMV RNA3 in which *MP* gene was exchanged by the indicated plant 30K-like endogenous genes. Schematic representation shows the GFP/AMV/CP (1) and the AMV RNA 3 derivatives containing the endogenous MPs from four different plant species (2–5) plus a negative control (6); see section 2.6 for details. Reading frames encoding the GFP, MP and coat protein (CP) are represented by green, red and yellow boxes, respectively. The C-terminal 44 amino acids of the AMV MP are indicated as “A44”. The numbers in the boxes represent the total amino acids residues of the corresponding MP. The *Nco*I and *Nhe*I restriction sites used to exchange the MP gene are indicated. Images at the right correspond to representative pictures of the size of infection foci observed on inoculated P12 leaves. Fluorescence was monitored with a confocal laser scanning microscope at 3 dpi. The bars in the pictures represents 200 µm. The LaPlace estimator of the percentages of infection foci *versus* the single infected cells are indicated (±1 SD).

Henceforth, here we conclude that endogenous MPs may still retain their ancestral function as cell-to-cell movement, though not in a very efficient manner, in some plant species. Variation in this activity may also exist among plant species, although our limited sample size prevents making generalizations.

## 4. Discussion

Here we report results from preliminary experiments aiming to characterize the functional role of a plant NIRV with strong similarity with the caulimovirus MP from the 30K-superfamily that was recently found widespread in the genome of vascular plants (Mushegian and Elena, 2015). Firstly, we have collected preliminary data suggesting that some of these NIRVs may still be active to translocate RNAs or ribonucleoproteins from cell to cell via the plasmodesmata, at least partially. Previous results obtained with the MP of CaMV showed that this protein is fully functional in the AMV context (Sánchez-Navarro et al., 2010), allowing the cell-to-cell and systemic transport of AMV. However, the limited viral transport observed with the analyzed NIRVs points to the idea that the integrated viral proteins may have been exaptated and play some other role, yet to be well defined. Secondly, we have generated RNAi-based knock-down mutants of this gene in *A. thaliana* and sought for possible phenotypes. Identifying the phenotype of knockdown mutations is not always an easy task. Here we have explored differences in development between the mutants and the wildtype plants, as well as for differences in resistance to abiotic (*e.g*., salinity) and biotic (*e.g*., response to infection with a panel of viruses) stresses. Our results suggest that knocking-down the expression of Arabidopsis’ endogenous 30K-like *AtMP* results in a small delay in plant development, a reduction of germination rate that feeds-back with an increase in sensitivity to salinity, and an enhancement in resistance to at least one RNA virus. Future work will further explore these observations.

Horizontal gene transfer from RNA virus genomes into the genome of their eukaryotic hosts has a tremendous evolutionary potential both for the cell and for the virus. As already illustrated in the Introduction, cells may acquire new functions from EVEs by different mechanism, including mutagenesis and the induction of genomic rearrangements, alterations of gene expression (reviewed in Feschotte and Gilbert, 2012), the exaptation of viral functions (Taylor and Bruenn, 2009; Katzourakis and Gifford, 2010), or most notably from an infectious disease perspective, acquiring resistances to viral pathogens by a sort of integration-based immunity (*e.g*., Bejarano et al., 1996; Bertsch et al., 2009; Taylor and Bruenn, 2009). Indeed, many successful resistances against plant viruses are based in the transgenic expression of viral proteins. Illustrative examples being tobacco plants expressing the coat protein of PVY (Van der Vlugt et al., 1992) or the replicase protein of TMV (Golemboski et al., 1990), tomatoes expressing the BV_1_ or BC_1_ proteins from bean dwarf mosaic begomovirus (Hou et al., 2000), or papayas expressing the coat protein the papaya ringspot potyvirus (Gonsalves, 1998). Although in the early times of this biotechnological application the molecular mechanisms were not understood, later on it was shown that resistance was mediated by the production of siRNAs during the transcription of these transgenes (Hamilton and Baulcombe, 1999; Tenllado et al., 2004).

A good effort has gone towards understanding and illustrating the origin of EVEs and their role in hosts evolution. However, the effect of EVEs in populations of viruses carrying homologous sequences has not received much attention so far. Viral transgenes can be considered as a particular case of EVEs, and therefore, transgenic expression of viral genes by plants may provide valuable information to cover this gap. One can envision situations in which the expression of a transgene is not efficiently inducing RNAi responses or the siRNAs are not homogeneously distributed in the plant, or situations in which virus carry strong suppressors of RNA silencing that allow for certain level of viral replication and escape from the surveillance of AGO complexes, or even situations of coinfection with other viruses that generate positive synergisms (Martínez et al., 2013). Under such situations, it is even possible to imagine that the product of the transgene may even complement defective viral genomes. This was, indeed, the case of transgenic plants expressing a fragment of the CP of cowpea chlorotic mottle bromovirus (CCMV) (Greene and Allison, 1994). When these transgenic plants were inoculated with an engineered genotype of CCMV that contained a deletion of the same CP region, systemic infections were observed and full-genome viruses recovered. Similar results have also been described for cucumber mosaic cucumovirus (Morroni et al., 2009, 2013), and the MPs of TMV and red clover necrotic mosaic dianthovirus (Giesman-Cookmeyer et al., 1995; Rao et al., 1998). Together, these studies show that recombination is likely to occur between mRNAs from transgenes of viral origin and the viral genome. Another illustrative example of the evolutionary effect of infecting a transgenic plant expressing one of the virus’ genes was provided by Tromas et al. (2014). In this case, tobacco plants expressing the NIb protein of tobacco etch potyvirus (TEV) were infected with a wildtype genotype of TEV. NIb is the replicase of TEV, thus meaning that any individual virus genome containing a deletion of a part or the entire gene would necessarily be non-viable by itself. Indeed, this was proved by constructing a TEV-Δ*NIb* genotype: this virus was not viable in wildtype tobaccos. By contrast, it was fully viable in the transgenic plants, showing that the transgene product complemented for the viral defect (Li and Carrington, 1995; Tromas et al., 2014). An evolution experiment was set up, allowing multiple independent lineages of TEV to evolve in transgenic plants. Of relevance for the discussion here, some of the evolving lineages accumulated genomes containing different deletions in the *NIb* gene. Most lineages perfectly removed the entire *NIb*, while others lacked pieces of it. Interestingly, when TEV-Δ*NIb* was competed against TEV in the transgenic plants, the former won the competition, thus showing it has a fitness advantage. Whether this advantage was owed to its shorter genome and faster replication or to the escape of siRNAs, was not tested. In any case, the study nicely illustrates two points: (*i*) as extreme cellular parasites they are, RNA viruses can utilize any resource provided by the cell; the gene products from NIRVs may be one of such resources. (*ii*) By doing so, RNA viruses may streamline even further their genomes.

## Acknowledgement

We thank Arcady Mushegian for motivating us to do this study and Francisca de la Iglesia, Paula Agudo and Lorena Corachan for excellent technical assistance. This research was supported by grants from Spain’s Agencia Estatal de Investigación - FEDER (BFU2015-65037-P and BIO2014-54862-R) and Generalitat Valenciana (PROMETEOII/2014/021 and PROMETEOII/2015/010).

## Appendix A. Supplementary data

Supplementary data associated with this article can be found, in the online version, at X.

---

**Table S1.**
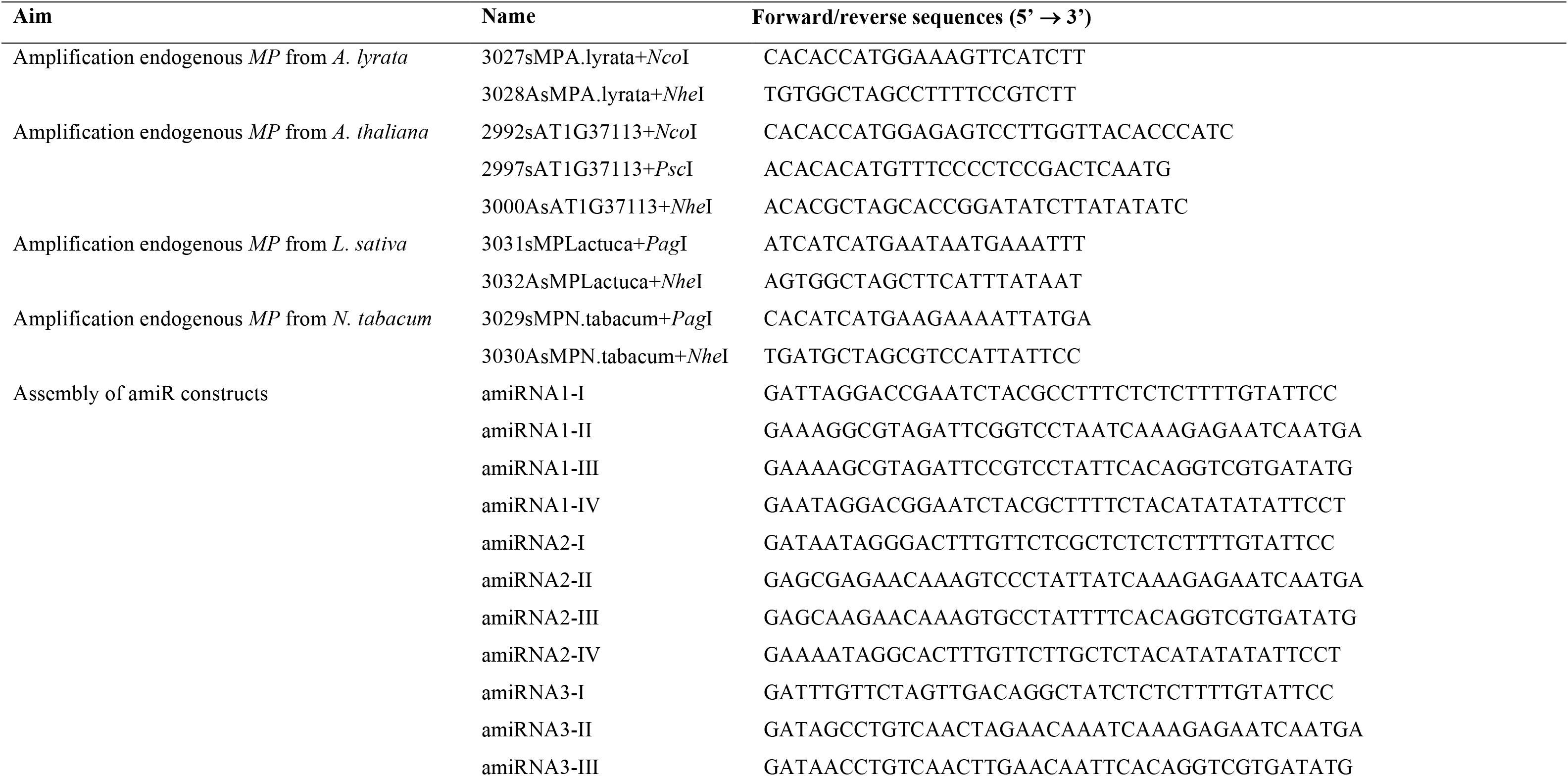
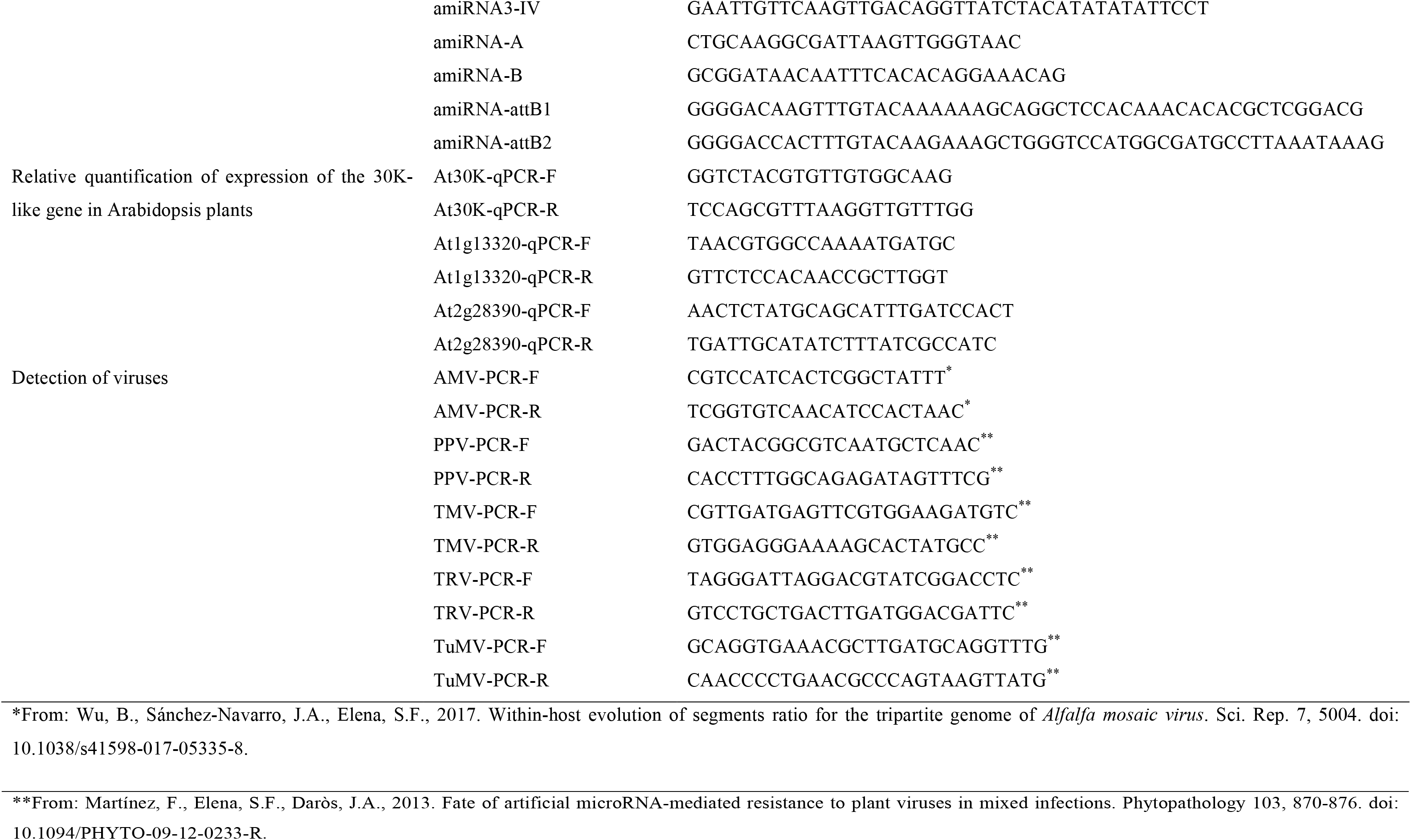
List of primers used in this study.

